# Transcriptome-based design of antisense inhibitors re-sensitizes CRE *E. coli* to carbapenems

**DOI:** 10.1101/2019.12.16.878389

**Authors:** Thomas R. Aunins, Keesha E. Erickson, Anushree Chatterjee

## Abstract

Carbapenems are a powerful class of antibiotics, often used as a last-resort treatment to eradicate multidrug-resistant infections. In recent years, however, the incidence of carbapenem-resistant *Enterobacteriaceae* (CRE) has risen substantially, and the study of bacterial resistance mechanisms has become increasingly important for antibiotic development. Although much research has focused on genomic contributors to carbapenem resistance, relatively few studies have examined CRE pathogens through changes in gene expression. In this research, we used transcriptomics to study a CRE *Escherichia coli* clinical isolate that is sensitive to meropenem but resistant to ertapenem, to both explore carbapenem resistance and identify novel gene knockdown targets for carbapenem re-sensitization. We sequenced total and small RNA to analyze gene expression changes in response to treatment with ertapenem or meropenem, as compared to an untreated control. Significant expression changes were found in genes related to motility, maltodextrin metabolism, the formate hydrogenlyase complex, and the general stress response. To validate these transcriptomic findings, we used our lab’s Facile Accelerated Specific Therapeutic (FAST) platform to create antisense peptide nucleic acids (PNA), gene-specific molecules designed to inhibit protein translation. FAST PNA were designed to inhibit the pathways identified in our transcriptomic analysis, and each PNA was then tested in combination with each carbapenem to assess its effect on the antibiotics’ minimum inhibitory concentrations. We observed significant treatment interaction with five different PNAs across six PNA-antibiotic combinations. Inhibition of the genes *hycA*, *dsrB*, and *bolA* were found to re-sensitize CRE *E. coli* to carbapenems, whereas inhibition of the genes *flhC* and *ygaC* was found to confer added resistance. Our results identify new resistance factors that are regulated at the level of gene expression, and demonstrate for the first time that transcriptomic analysis is a potent tool for designing antibiotic PNA.

## Introduction

In recent years, the emergence of carbapenem-resistant *Enterobacteriaceae* (CRE) has been marked by the World Health Organization (WHO) as a critical priority for antibiotic development.^1^ Resistance to carbapenems, a subclass of the cell wall-targeting β-lactams that are often termed antibiotics of last resort,^2^ has become increasingly common in recent decades,^3–5^ resulting in rising threats of infection mortality.^6–9^ In 2017, the Centers for Disease Control estimated that, in the U.S., 13,100 infections and 1,100 deaths per year were caused by carbapenem-resistant *Enterobacteriaceae* alone.^10^ Furthermore, the WHO has called for further investment in both basic research and clinical development for CRE pathogens, as the current pipeline is not sufficient to address the threat.^11^ Our study seeks to use transcriptomics both to better understand carbapenem resistance and to design new methods to combat its development. Each of these aims is facilitated by the application of our Facile Accelerated Specific Therapeutic (FAST) platform, a semi-automated pipeline for the design, synthesis, and testing of antisense peptide nucleic acids (PNA)—synthetic DNA analogues with peptide backbones that can inhibit translation on a gene-specific basis. Here, FAST PNA allow us to validate our conclusions from our transcriptomic analysis, and to restore carbapenem susceptibility to a CRE clinical isolate.

Structural modifications, particularly a *trans*-α-1-hydroxylethyl substituent at position 6, are believed to endow carbapenems with higher stability against β-lactamases,^12–14^ as well as broader observed efficacy,^15, 16^ compared with other β-lactam drugs. However, increasing prevalence of carbapenemase enzymes in *Enterobacteriaceae*—such as oxacillinase-48, metallo-β-lactamases, and *Klebsiella pneumoniae* carbapenemases—provide mechanisms for the development of resistance.^3, 5, 17, 18^ Multiple studies have found that pathogenic bacteria without carbapenemases can acquire carbapenem resistance through the combination of β-lactamases and outer membrane porin deficiencies (neither factor alone was sufficient for resistance),^19–23^ through the expression of low-affinity or mutated penicillin-binding proteins,^24–27^ or through the overproduction of efflux pumps.^28, 29^

Other research has sought to understand resistance by tracking transcriptomic changes in carbapenem-resistant pathogens. Two studies on the gene expression profile of carbapenem-resistant *Acinetobacter baumanii* found significant upregulation of transposable elements, recombinase, and other mutation-encouraging factors, in addition to the expected upregulation of efflux pump and β-lactamase genes.^30, 31^ Four recent studies that examined transcriptomic profiles of CRE*—Enterobacter cloacae*, *K. pneumoniae* (2), and *Escherichia coli*—identified less consistent responses, although downregulation of porin genes and upregulation of cell survival and β-lactamase genes were observed.^32–35^ Of this prior research, only one study^32^ attempts to evaluate the transcriptomic response of a resistant *Enterobacteriaceae* strain under antibiotic challenge, the majority examining only the constitutive gene expression levels of an untreated resistant pathogen.

In this study, we both explore and counter carbapenem resistance by examining the short-term (<1 hr) transient transcriptomic response of a clinical isolate of multidrug resistant (MDR) *E. coli* (referred to from here as *E. coli* CUS2B) to carbapenem treatment. The strain demonstrates resistance to ertapenem, but sensitivity to meropenem and doripenem, and we sought to use this partial carbapenem resistance phenotype to help understand the development of resistance in *Enterobacteriaceae*. We used a combination of total and small RNA sequencing to assess the response of *E. coli* CUS2B to either ertapenem or meropenem treatment and identify genes that were potentially important to the pathogen’s resistance profile. We then used these genes, together with genome sequencing of the clinical isolate, as inputs for the FAST platform.^36^ For each target gene, we designed and synthesized PNA molecules that were predicted to inhibit translation via sequence-specific binding to the mRNA translation initiation region. Growth assays of PNA-carbapenem combination treatments were used to validate each gene’s relevance to the resistance phenotype, and to determine whether PNA designed using transcriptomics could re-sensitize *E. coli* CUS2B to sub-MIC carbapenem concentrations.

## Results

### E. coli CUS2B: a multidrug-resistant Enterobacteriaceae with partial carbapenem resistance

To validate the *E. coli* CUS2B resistance phenotype observed in the clinic, we measured the isolate’s minimum inhibitory concentrations (MIC) for a variety of antibiotics from different classes (Fig. 1A). We found that *E. coli* CUS2B was resistant to almost all antibiotics (based on breakpoints defined by the Clinical & Laboratory Standards Institute^37^), including multiple penicillins and ertapenem. Two potent carbapenem antibiotics, meropenem and doripenem, were the only drugs to which the clinical isolate was susceptible. To investigate this partial carbapenem resistance, we focused on the *E. coli* CUS2B response to ertapenem and meropenem (Fig. 1B). The structures of meropenem and ertapenem differ in the pyrrolidinyl ring’s position 2 side chain (Fig. 1C). Meropenem’s substituent amide group is thought to be responsible for increased potency against gram-negative organisms in comparison to imipenem.^38^ At this position ertapenem has a benzoate substituent group, which imbues the molecule with a net negative charge and increases its lipophilicity, resulting in increased plasma half-life but decreased affinity for membrane porins.^13^

**Figure 1.**
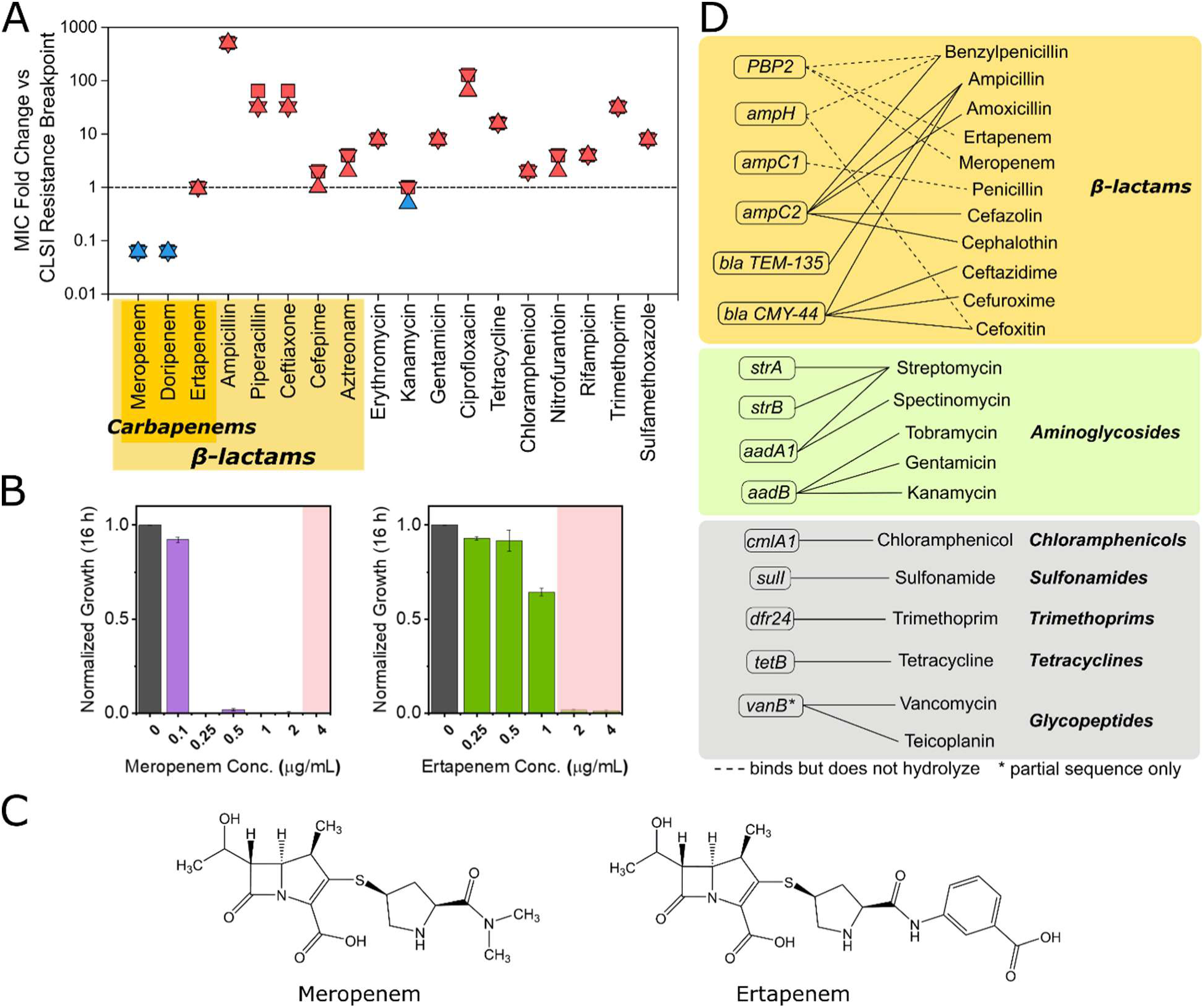
Resistance profile and resistance genes of *E. coli* CUS2B. (A) Minimum inhibitory concentration (MIC) of *E. coli* CUS2B for 18 antibiotics. The MIC is shown as the fold change in MIC vs. the CLSI resistance breakpoint. Values ≥ 1 indicate resistance, while values < 1 indicate susceptibility. (B) Normalized 16-hour endpoint optical density (590 nm) for a range of concentrations of meropenem or ertapenem. The resistance breakpoint is highlighted in pink for each antibiotic. (C) Structure of meropenem and ertapenem. (D) Antibiotic resistance genes identified in CU2SB from whole-genome sequencing data, and antibiotics to which the genes are predicted to confer resistance.

### Resistance factor identification via whole-genome sequencing

We performed whole genome shotgun sequencing for two purposes: (1) to create a genome assembly that could be used for antisense PNA design, and (2) to search for genomic contributions to the resistance phenotype. Using the ARG-ANNOT database,^39^ we found that the strain encodes fifteen genes related to antibiotic resistance (Fig 1D), including six associated with β-lactams either as penicillin binding proteins (PBPs) or β-lactamase enzymes. Four of these six—labeled by ARG-ANNOT as *ampC1*, *ampC2*, *ampH*, and a generic PBP—are encoded chromosomally, and have high-similarity homologues in *E. coli* reference strains (Table S1). The generic PBP, which was found to identically match the protein sequence for *E. coli* MG1655 *mrdA*/PBP2, is known to bind both carbapenems tested; PBP2 is the PBP to which meropenem and ertapenem demonstrate greatest activity.^18, 40, 41^ The genes *ampC1* and *ampH* show >96% homology with *E. coli* MG1655 PBPs *yfeW*/PBP4B and *ampH*/AmpH, respectively, and neither has been found to bind with carbapenems. The gene *ampC2* shows 98% nucleotide homology with *E. coli* MG1655 β-lactamase *ampC*/AmpC, but has ten altered codons from the reference sequence (Table S1). Additionally, the −35 to −10 promoter region for the CUS2B *ampC* has four mismatched nucleotides from the same promoter region in MG1655, which may alter its expression level relative to the basal non-resistant levels in the reference strain. The remaining two β-lactam-associated genes are plasmid encoded; these are the β-lactamases TEM-135 and CMY-44. *E. coli* CUS2B also encodes the outer membrane porins OmpA, OmpC, and OmpF, the mutation or downregulation of which may influence carbapenem efficacy.^19, 33, 42^ These three proteins have, respectively, 95%, 90%, and 90% nucleotide homology with the corresponding genes in *E. coli* MG1655.

**Table 1.** Summary of combination treatment significant interactions, with normalized S-values (see Methods) calculated relative to drug combination predictions from the Bliss Independence Model. Plus/minus values represent standard error of the *S*-value. Bolded/colored numbers indicate P < 0.05/synergy (red) or antagonism (blue), after adjustment for multiple hypothesis testing.

| *PNA* | 24 h Growth |  | 24 h Cell Viability |  |
| --- | --- | --- | --- | --- |
|  | *Ertapenem* | *Meropenem* | *Ertapenem* | *Meropenem* |
| α-*ampC* 10 μM | **0.76 ± 0.11** | -0.06 ± 0.18 | **0.44 ± 0.22** | **-0.2 ± 0.09** |
| α-*hycA* 10 μM | **0.23 ± 0.06** | -- | 0.36 ± 0.20 | -- |
| α-*dsrB* 10 μM | **0.85 ± 0.06** | -- | **0.73 ± 0.20** | -- |
| α-*bolA* 10 μM | **0.83 ± 0.02** | -- | **0.75 ± 0.09** | -- |
| α-*bolA* 15 μM | -- | **0.61 ± 0.13** | -- | **0.77 ± 0.12** |
| α-*flhC* 10 μM | -- | **-0.90 ± 0.02** | -- | **-1.11 ± 0.08** |
| α-*ygaC* 10 μM | -- | **-0.93 ± 0.00** | -- | **-0.90 ± 0.04** |

### Profiling gene expression in response to ertapenem and meropenem treatment

In previous work, we have established the ability of PNA and the FAST platform to design effective antibiotic potentiators through the use of genomic data, using either knockouts of essential genes or resistance genes.^36, 43^ Here, FAST PNA allow us to explore transcriptomic analysis and determine whether it can be similarly useful in reversing a bacterial resistance phenotype. First, to reveal possible non-genetic contributions to the strain’s carbapenem resistance profile, we exposed exponentially growing *E. coli* CUS2B to ertapenem and meropenem and examined gene expression profiles after thirty and sixty minutes of treatment (Fig. 2A). This exposure time was selected based on observations from others that 30-60 minutes is favorable for locating gene expression changes specific to antibiotic exposure.^44–46^ The short time frame also greatly reduces the likelihood that the gene expression signal will be confounded by the emergence of one or more mutant genotypes. *E. coli* CUS2B was diluted 1:20 from overnight cultures and grown for 1 hour to exponential phase prior to treatment with 2 µg/mL of ertapenem or 1 µg/mL of meropenem. Each carbapenem concentration is half of that required to eradicate *E. coli* CUS2B under these culture conditions (conditions differ from Fig. 1 MIC assay) (Fig. S1).

**Figure 2.**
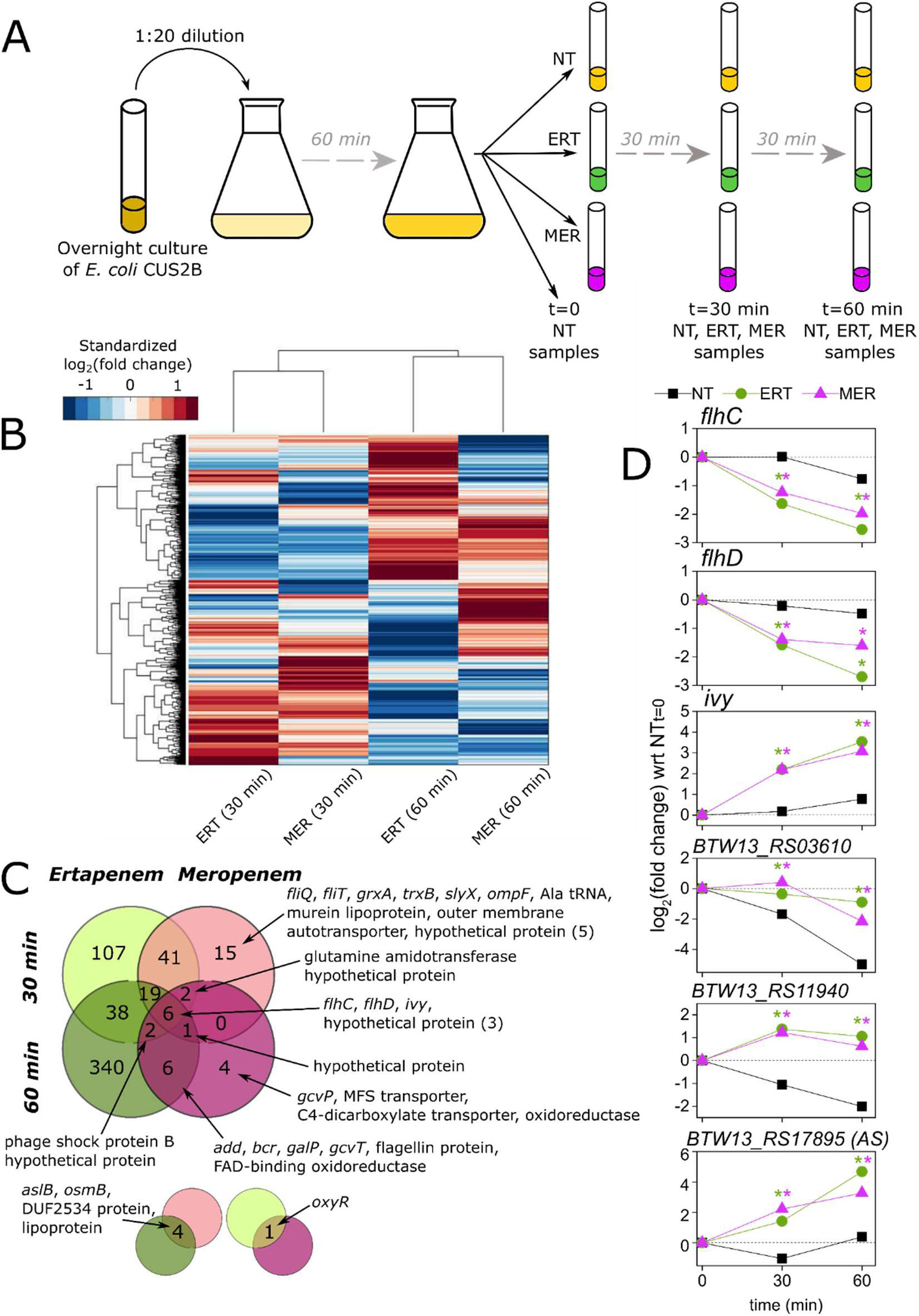
Total RNA-sequencing uncovers ertapenem response and identifies motility genes as resistance factors. (A) Overview of the sample collection protocol for RNA sequencing. Exponential phase cells were collected after 0, 30, and 60 minutes of growth in either CAMHB with no antibiotic (NT), meropenem (MER), or ertapenem (ERT). (B) Hierarchical clustering of gene expression values in ertapenem and meropenem-treated *E. coli* CUS2B. Values are log2(fold change), with respect to the corresponding no treatment duplicates at the same timepoint. For visual clarity, the heatmap has been standardized such that the mean is 0 and the SD is 1 across each gene (each row). (C) Intersections in significantly differentially expressed (DE) genes between stress conditions and timepoints. (D) Time course of gene expression changes for the 6 transcripts that were DE in ertapenem and meropenem at both 30 and 60 minutes. Asterisks indicate significant DE (*P* < 0.05) vs. the no treatment condition at the same timepoint. NT = no treatment, ERT = ertapenem, MER = meropenem.

We identified differentially expressed (DE) genes by comparing the RNA sequencing data from ertapenem- and meropenem-treated samples to an untreated control at the same timepoint. The DESeq R package^47^ was used to evaluate significance and correct *P*-values for multiple hypothesis testing. General expression trends were evaluated using hierarchical clustering across genes and conditions (Fig. 2B). Conditions were found to cluster by timepoint rather than antibiotic—an effect previously observed in a transcriptomic study of vancomycin-challenged *Staphylococcus aureus*^48^*—*which suggests a generalized and transient antibiotic response. In our analysis, we detected 41 transcripts that were DE in both treatments after 30 minutes of exposure, six transcripts DE in both antibiotics after 60 minutes of exposure, and six that were DE in both treatments at 30 and 60 minutes (Fig. 2C, File S1).

The six DE genes common to all conditions are indicated in Fig. 2D. Two of these genes, *flhC* and *flhD*, code for components of the transcriptional regulator FlhDC, which is responsible for regulating motility-associated functions such as swarming and flagellum biosynthesis.^49^ Both *flhC* and *flhD* genes were significantly underexpressed at 30 and 60 minutes. Perhaps relatedly, motility-associated gene ontology terms (GO:0040011: locomotion; GO:0071918: bacterial-type flagellum dependent swarming motility) were significantly overrepresented^50^ within the 30-minute overlapping set, accounting for 26 of the total 41 genes (Table S2). All 26 were underexpressed. This effect was diminished by 60 minutes, with only *flhC*, *flhD*, and *fliC* (the gene encoding flagellin^51^) remaining significantly underexpressed.

The gene *ivy*, an inhibitor of bactericidal vertebrate lysozymes,^52, 53^ was overexpressed in both treatments at 30 and 60 minutes. Both lysozymes and carbapenems disrupt peptidoglycan polymerization, although *ivy* is not known to interact with these antibiotics. Three other transcripts of unknown function were overexpressed in all conditions: BTW13_RS03610 (*ymgD* superfamily), BTW13_RS11940 (DUF1176 superfamily), and a transcript antisense to BTW13_RS17895 (putative lipoprotein, DUF1615 superfamily).

In the ertapenem response we find many more DE genes than in the meropenem response, including 38 DE genes shared between the two time points (compared with none shared across both meropenem time points). Within this set, we observed significant overrepresentation of genes related to maltodextrin transport (*mal* operon, GO:0042956) and the ferredoxin hydrogenase complex (*hyc* operon, GO:0009375).^50^ All of these overrepresented genes were found to be overexpressed in ertapenem treatments. Only three genes were underexpressed in ertapenem at both 30 and 60 minutes: *lptG*, a member of the lipopolysaccharide transport system, *phoH*, an ATP-binding protein, and *cstA*, a starvation-induced peptide transporter. Of the genes specific to the meropenem response, only the flagellar biosynthesis proteins *fliQ* and *fliT* are related, and the downregulation of these genes did not continue to the 60-minute timepoint.

We also searched for differential expression in outer membrane porin operon (*omp*) genes, previously linked to carbapenem-resistance,^19, 33, 42^ and resistance-related genes identified by ARG-ANNOT. Of the *omp* operon, only *ompF* was found to be significantly DE in any condition with respect to no treatment (underexpressed in meropenem, 30 minutes) (Fig. S2), while *ompA* and *ompC* expression tracked closely with the no treatment conditions in all experiments. When expression levels of the ertapenem and meropenem experiments were directly compared at each time point, none of the three genes were found to be significantly DE. No resistance-related genes were DE in any condition (Fig. S3, File S1).

Based on these observations, we chose three genes to target using PNA: *hycA*, *malT*, and *flhC*. The former two genes were chosen to probe the *hyc* and *mal* operons for their importance to resistance and their utility as antibiotic re-sensitization targets. The gene *flhC* was chosen to validate the consistent downregulation of the FlhDC system and evaluate whether further knockout of the gene would confer greater carbapenem resistance.

### Differential expression of small RNA

In addition to total RNA sequencing, we performed small RNA sequencing to search for resistance contributions and potential FAST PNA targets among short nucleic acids potentially involved in gene regulation. Small RNA have been previously shown to influence bacterial stress and antibiotic response.^54–56^ We used an RNA isolation protocol that enriched for sRNA (see Methods) prior to sequencing, and sequencing data were aligned to the *E. coli* UMN026 genome (the reference that maximized alignment homology) using the Rockhopper^57^ pipeline, which allowed for identification of previously documented sRNA and novel RNAs.

We observe more overlap of DE genes between single time points than between respective antibiotic treatments, suggesting a generalized and transient response similar to that of total RNA expression (Fig. 3A). We find 22 sRNAs to be DE in at least two out of the four conditions (Fig. 3B), including known regulatory sRNAs (*dicF*, *ssrA*), annotated short protein-coding genes (*ilvB*, *acpP*, *bolA*, *csrA*, *ihfA*, *lspA*), small putative protein-coding genes (*dsrB*, *yahM*, *ybcJ*, *ygdI*, *ygdR*, *ytfK*), small transcripts antisense to coding genes (*ygaC*, *hemN*, ECUMN_1534/5), and novel predicted transcripts.

**Figure 3.**
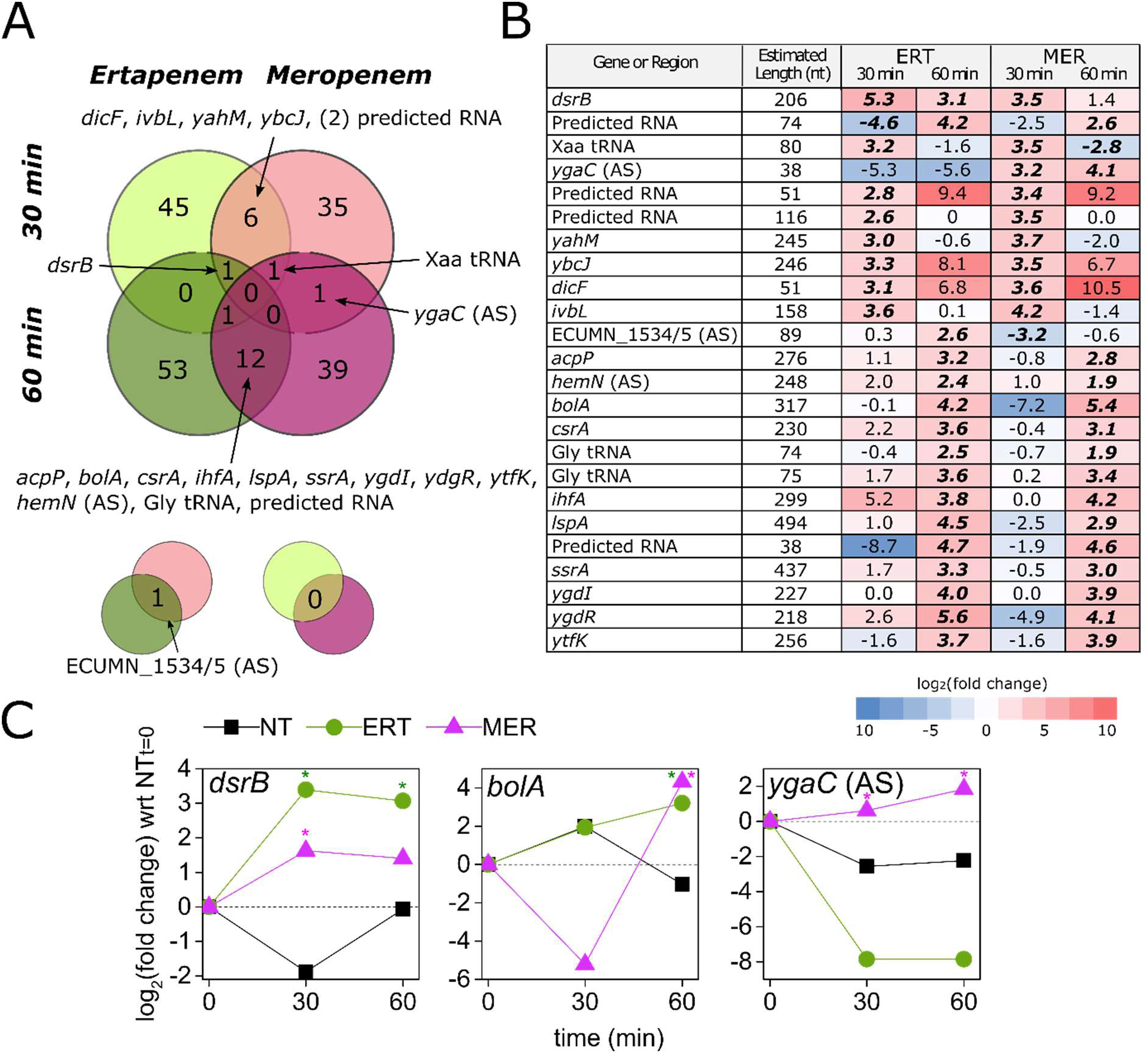
Small RNA-sequencing identifies regulator genes and new targets for PNA antibiotics. (A) Overlaps between conditions in significantly differentially expressed (DE) transcripts from the small RNA enriched pool. (B) Detail on the 22 RNAs that were DE in at least two conditions. Log_2_(fold change) values here are with respect to the no treatment condition from the same timepoint. Values are bold and italic if *P* < 0.05 in that condition. (C) Time course of gene expression for four small RNA transcripts of interest. Log_2_(fold change) here is with respect to the t=0 min samples. Asterisks indicate *P* < 0.05, with comparison to the no treatment condition from the same timepoint. NT = no treatment, ERT = ertapenem, MER = meropenem

From these lists we chose three genes to investigate with FAST PNA: *bolA*, *dsrB*, and *ygaC*. *bolA* was chosen based on its relation to PBPs, AmpC,^58^ and the cellular stress response,^59^ whereas the latter two were chosen to discriminate between the two carbapenem responses. The time course of gene expression for three small transcripts of interest is presented in Fig. 3C. We found *bolA* to be overexpressed in meropenem and ertapenem at 60 minutes, which may indicate that both antibiotics are being detected and able to activate *bolA*, but the subsequent response is only effective against ertapenem. *dsrB* was overexpressed in ertapenem at both time points, but in meropenem only at 30 minutes. Although the function of *dsrB* is unknown, it is controlled by σ^S^, the general stress response and stationary-phase sigma factor.^60^ The transcript antisense to *ygaC* was overexpressed in meropenem at both timepoints but was not detected in ertapenem-treated populations. The function of *ygaC* is unknown, but it is controlled by the Fur transcriptional dual regulator, which suggests that it may have a role in stress response.

### PNA antisense inhibition of RNA sequencing targets

The FAST platform comprises Design, Build, and Test modules for the creation of antisense PNA (Fig. 4A). In this study, we began our design process using transcriptomic data to generate a list of target genes, which, together with a whole-genome assembly and genome annotation, were used as inputs for the FAST tool PNA Finder. This tool was used to design multiple antisense PNA candidates for each gene target, with 12-mer sequences—a length that seeks to optimize both specificity^61–63^ and transmembrane transport^64^—that were complementary to mRNA nucleotide sequences surrounding the translation start codon. PNA Finder then filtered this set of candidates to minimize the number of predicted off-targets within the *E. coli* CUS2B genome, to maximize solubility,^65^ and to avoid any self-complementing sequences. For the FAST Build module, a single PNA for each gene target was selected and synthesized using Fmoc chemistry, with the cell-penetrating peptide (KFF)_3_K attached at the N-terminus to improve transport across the bacterial membrane. These PNA were then tested in *E. coli* CUS2B cultures in combination with each carbapenem to determine whether the two treatments would interact as predicted. A two-way ANOVA test was used to assess interaction significance, and normalized *S*-values (see Methods) were used to compare the observed growth to the expected growth, as predicted by the Bliss Independence Model for drug combinations.^66^

**Figure 4.**
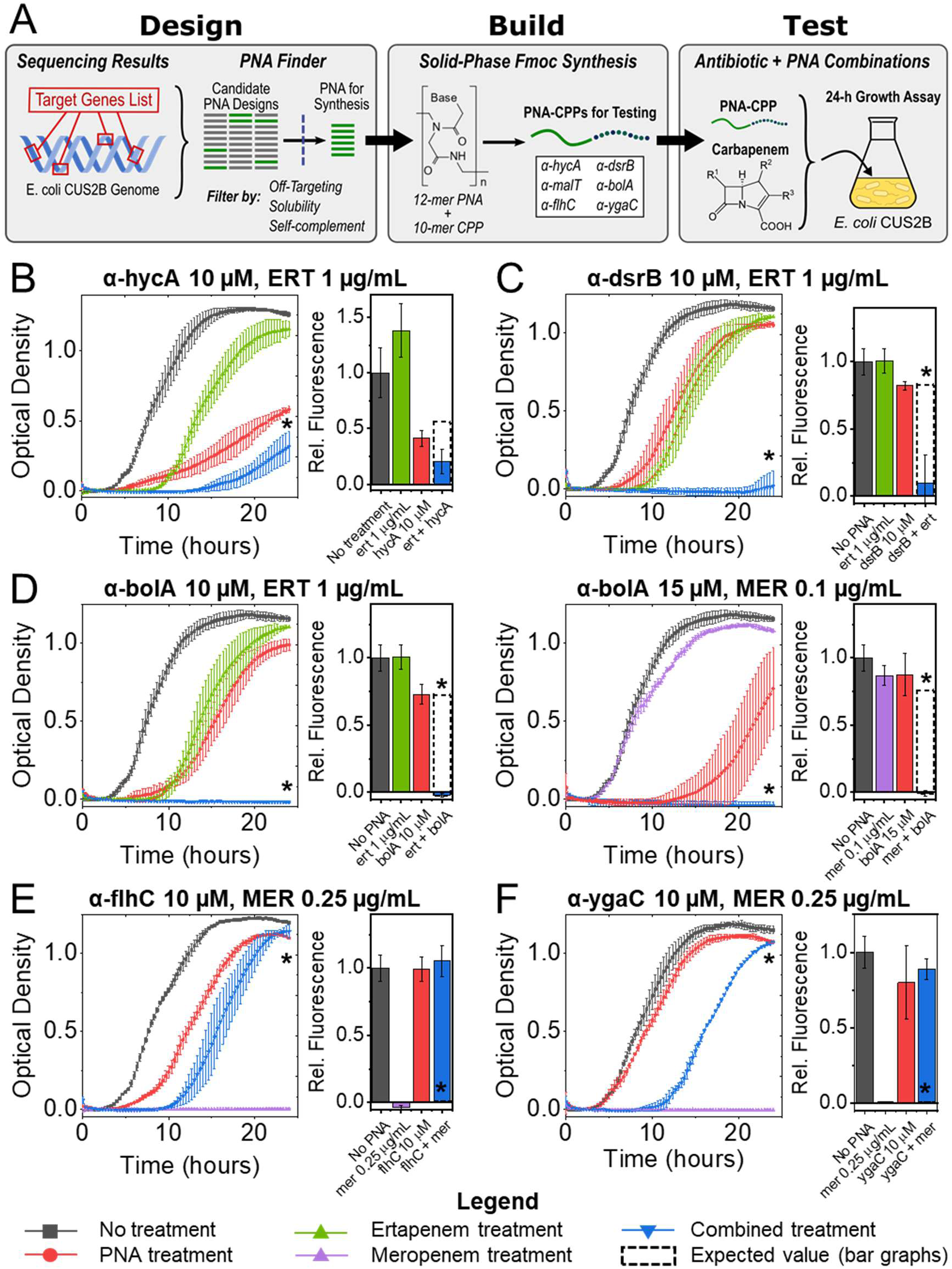
Application of the FAST platform to the gene targets identified in transcriptomic analysis. (A) Schematic of the FAST platform’s Design, Build, and Test modules, as they were applied to the conclusions generated by our gene expression experiments. (B-F) Growth curves and endpoint cell viability assays for CRE *E. coli* PNA-antibiotic combinations. Curves are shown for experiments in which significant interaction was observed: (B) α-hycA (10 µM) combined with ertapenem (1 µg/mL); (C) α-dsrB (10 µM) combined with ertapenem (1 µg/mL); (D) α-bolA (10 µM) combined with ertapenem (1 µg/mL), and α-bolA (15 µM) combined with meropenem (0.1 µg/mL); (E) α-flhC (10 µM) combined with meropenem (0.25 µg/mL); (F) α-ygaC (10 µM) combined with meropenem (0.25 µg/mL). ERT = ertapenem, MER = meropenem. *: ANOVA interaction P-value < 0.05, corrected for multiple hypothesis testing via the Benjamini-Hochberg procedure.

Based on the results of our transcriptomic analysis, we selected three genes identified by our total RNA sequencing analysis (*hycA*, *malT*, and *flhC*) and three genes identified by our small RNA sequencing analysis (*bolA*, *dsrB*, and *ygaC*) to be targeted by FAST PNA. Differential expression of *flhC* and *bolA* was observed in both carbapenems, while differential expression of *hycA*, *dsrB*, and the operon controlled by *malT* was prevalent in the ertapenem response. The transcript antisense to *ygaC* was overexpressed in both meropenem conditions, but the gene itself was not differentially expressed. While we suspect that the *ygaC* antisense transcript regulates the gene *ygaC*, PNA have not been established to interfere with ncRNA regulation. With this in mind, we designed a PNA to bind to the *ygaC* sense transcript and inhibit protein translation, to test the hypothesis that *ygaC* inhibition may confer greater meropenem resistance in a similar manner to the *flhC*-targeted PNA. The genome assembly for the clinical isolate was used by FAST to design multiple PNA for each selected gene, which were then screened for high solubility, minimal self-complementarity, and zero off-target gene inhibition in *E. coli* CUS2B (Table S3). Additionally, a scrambled-sequence nonsense PNA was designed to control for any possible effects of the PNA or CPP independent of sequence. No effects were found with nonsense PNA alone or in any nonsense PNA-antibiotic combination treatment (Fig. S5).

We also designed a PNA to inhibit the translation of the chromosomal β-lactamase AmpC, based on prior research showing that the enzyme can elevate ertapenem MICs.^67, 68^ This PNA was also synthesized to assess the relative effectiveness of PNA targets selected using transcriptomic analysis, in comparison to those selected on a genomic basis. We have previously shown the ability to re-sensitize MDR bacteria by targeting β-lactamases.^43^ In a combination of *α-ampC* (10 µM) in combination with ertapenem (1 µg/mL), we also observe significant synergistic interaction, evidenced both by the growth curve endpoint and cell viability assay (Table 1, Fig. S6A). *E. coli* CUS2B cultures treated with a combination of 0.1 µg/mL meropenem with 10 µM *α-ampC* grew similarly to cultures treated with meropenem alone, as expected. Although the comparison of treatments in the cell viability assay did demonstrate significant interaction, the data does not seem to indicate a combinatorial effect, as the fluorescence level is virtually unchanged across the three treated conditions. Additionally, increasing the concentration of *α-ampC* to 15 µM could not resolve any effect, as the PNA alone was virtually lethal at this concentration (Fig. S6B).

To determine whether transcriptomic analysis could be used by FAST to produce similar re-sensitization effects, *E. coli* CUS2B was treated with each of the six PNA at a concentration of 10 µM in combination with sub-MIC carbapenem treatments (1 µg/mL ertapenem, 0.1 µg/mL meropenem). We analyzed the cultures’ endpoint optical densities and cell viabilities (via resazurin assay) to assess interaction between the two treatments, based on a comparison with each individual PNA and carbapenem treatment. At these concentrations of PNA and antibiotic, we observed significant synergy between the PNAs *α-hycA*, *α-dsrB*, and *α-bolA* and ertapenem, with *S*-values of 0.23, 0.85 and 0.83 for their respective endpoint optical densities (Table 1, Fig. 4B-D). With each of these three PNAs, we performed additional interaction experiments at a PNA concentration of 15 µM with meropenem treatment to determine whether increased inhibition of these genes would demonstrate significant interaction. Of these combinations, 15 µM *α-bolA* demonstrated significant synergistic interaction (*S* = 0.61) with meropenem (Table 1, Fig. 4D). The PNA *α-ygaC*, *α-malT*, and *α-flhC* at concentrations of 10 µM did not exhibit significant interaction with ertapenem at 1 µg/mL or meropenem at 0.1 µg/mL (Fig. S7, Fig. S8A).

As noted above, we hypothesized that a combination of the PNA *α-flhC* or *α-ygaC* with carbapenem treatment would result in a recovery of growth and increased resistance. However, in the PNA-carbapenem combination treatments we did not observe growth recovery relative to the carbapenem-only treatment for the sub-MIC concentrations. We hypothesized that effects at such concentrations could be difficult to resolve, given that the growth curves of carbapenem-treated *E. coli* CUS2B reached endpoints similar to the untreated condition (Fig. S7A, Fig. S8A). To examine this hypothesis, we treated the clinical isolate with *α-flhC* or *α-ygaC* at 10 µM in combination with ertapenem or meropenem at their MICs (Fig. 1C; 2 µg/mL and 0.25 µg/mL, respectively). At these higher carbapenem concentrations, we observed significant antagonistic interaction between both PNA and meropenem in combination (*S* = –0.9 and *S* = –0.93, respectively, for endpoint optical density) (Table 1, Fig. 4E), whereas a combination of either PNA with ertapenem showed no growth rescue (Table 1, Fig. S8B).

## Discussion

In this study, we sought to combine transcriptomic analysis of carbapenem resistance with our FAST platform, both to better understand the partial carbapenem resistance profile of *E. coli* CUS2B, and to engineer carbapenem re-sensitization in the clinical isolate. Our results offer insights into pathways that are important for the development of resistance, identifying multiple, and demonstrate the first use of transcriptomics in the design of PNA for antibiotic applications.

To begin our analysis of the clinical isolate, we first used whole genome sequencing to search for any possible genomic resistance factors. We identified 15 genes related to antibiotic resistance, including six related to β-lactam antibiotics. None of these genes are dedicated carbapenemases, although the chromosomal *E. coli* AmpC β-lactamase is present. The basal expression of AmpC in wild-type *E. coli* is not enough to confer resistance to beta-lactams, but mutation-induced overproduction of the chromosomal AmpC in *Enterobacteriaceae* has been shown to promote antibiotic resistance.^33, 42, 69–71^ Furthermore, when coupled with porin deficiencies, AmpC expression can increase enterobacterial resistance to ertapenem while retaining susceptibility to other carbapenems.^23, 67, 68, 72, 73^ Our genomic analysis identified ten codon mutations in the *E. coli* CUS2B AmpC compared with *E. coli* MG1655, as well as four nucleotide mutations in the −35 to −10 promoter region. Furthermore, we identified numerous mutations of the *E. coli* CUS2B *omp* operon porin genes. It is possible that these mutations contribute to the ertapenem-resistant/meropenem-sensitive resistance profile that we observe in this clinical isolate. Using the FAST platform, we were able to design an anti-AmpC PNA to knockdown the translation of the β-lactamase and determine whether it has a significant role in resistance. As expected, we observed a strong significant synergistic combination between *α-ampC* and ertapenem, but not between *α-ampC* and meropenem. These observations agree with prior work from our lab that establish β-lactamases as viable re-sensitization targets, and serve as a basis of comparison for our transcriptomics-based PNA.

Transcriptomic analysis of carbapenem-challenged *E. coli* CUS2B revealed a much greater gene expression change in response to ertapenem than to meropenem: 485 DE genes were unique to ertapenem treatment versus 19 DE genes unique to meropenem. This may suggest the recognition of ertapenem and resultant response activation, in addition to innate basal resistance factors. In the DE analysis, we find no significant difference between the expression of outer membrane porin genes *ompA*, *ompC*, and *ompF* in the ertapenem experiments when compared to the meropenem experiments or the untreated conditions. Though *ompF* was found to be slightly underexpressed in meropenem after 30 minutes with respect to untreated conditions, the lack of differential expression in the ertapenem-treated conditions leads us to conclude that transcriptomic control of this gene is not a major resistance factor for *E. coli* CUS2B. As discussed above, the downregulation and deletion of these genes has previously been linked to carbapenem-resistance,^19, 33, 42, 67^ but, if this mechanism is active in *E. coli* CUS2B, it does not appear to be regulated by transient gene expression. Similarly, none of the resistance-related genes identified in our genomic analysis were found to be DE in any carbapenem treatment condition, and we may conclude that transcriptomic control of these genes does not contribute to the *E. coli* CUS2B resistance phenotype. Notably, though, this does not rule out a constitutively higher expression of the AmpC β-lactamase nor constitutively lower expression of porin genes, relative to the reference strain,

To determine the transient gene expression changes that contribute to the carbapenem resistance phenotype, we analyzed differential expression overlaps between the each of the four samples (two carbapenems, two time points each). The most apparent trend was an overwhelming underexpression of motility-related genes after 30 minutes of exposure to both antibiotics, an effect that lessened after 60 minutes of exposure. This response is consistent with prior work that has noted a decreased expression of motility genes in response to generalized environmental stressors.^74–77^ Two of these motility-related genes, *flhC* and *flhD*, were consistently DE in both treatments at both timepoints, along with the lysozyme inhibitor *ivy*, an RNA antisense to a putative lipoprotein, and two transcripts of unknown function.

We used antisense PNA to probe the resistance contributions of the *flhDC* operon. Interestingly, we observed significant antagonistic interaction between PNA inhibition of FlhC translation and meropenem treatment, but not ertapenem treatment. The deletion of the FlhDC regulator has been shown in *E. coli* to eliminate motility, and has been shown to cause downregulation of a large number of genes that are primarily related to chemotaxis/motility and flagellar surface structures.^74^ It is striking that this resistance factor can be artificially induced to so effectively rescue the *E. coli* CUS2B strain from carbapenem challenge, and that the effect is stronger than any response that the cell naturally activates, even over the course of 24 hours. Furthermore, the dissonance between transcriptomic results (underexpressed in all conditions) and PNA inhibition results shows that FlhC is downregulated as a general carbapenem response, but does not universally improve survival in resistant strains. These observations point to avenues by which enterobacteria may acquire greater degrees of carbapenem resistance.

We next examined the ertapenem-specific response in an effort to use transcriptomics to engineer carbapenem re-sensitization. We identified a total of 38 transcripts that were uniquely DE in ertapenem-treated samples. In this set we identified two operons—the *mal* operon (maltodextrin metabolism) and the *hyc* (formate hydrogenlyase complex) operon—that are consistently overexpressed. We designed and synthesized PNA inhibitors for *malT*, which, while not DE, is an activator of the overexpressed *mal* operon, as well as the DE gene *hycA*, in order to evaluate and interfere with the clinical isolate’s response to ertapenem. We observed significant synergistic growth inhibition between *α-hycA* and ertapenem, but not meropenem, while combinations of *α-malT* with the carbapenems did not produce significant treatment interaction.

The genes of the *hyc* operon code for formate hydrogenlyase complex, which mediates formate oxidation and has shown a potential connection to ATP synthesis.^78^ However, previous mutational analysis of *hycA* has provided evidence that the protein HycA works as a negative regulator of this system, as increased formate dehydrogenase activity was observed following its deletion.^79, 80^ Based on our observations, the upregulation of the formate hydrogenlyase complex is likely important to the *E. coli* CUS2B’s ertapenem resistance, but requires commensurate upregulation of the HycA regulator. The absence of this regulator results in the observed synergistically toxic effect when *α-hycA* is combined with ertapenem, an effect not observed in the meropenem response because the system is not activated by the antibiotic. These results validate the role of formate metabolism in the carbapenem response, and support the conclusion that transcriptomic data is valuable for engineering antibiotic re-sensitization in resistant enterobacteria.

In addition to applying FAST PNA to gene targets identified by total RNA sequencing, we also analyzed differential expression of small RNA in search of potential resistance factors. We identified 22 transcripts that were differentially expressed across one or more of the treatment conditions and timepoints, and selected three genes to be targeted by FAST: *dsrB*, *bolA*, and *ygaC*. *dsrB* and *bolA* are translated into short proteins, but no annotation for the *ygaC* antisense complement has been identified, leading to the hypothesis that the transcript serves as a regulatory RNA for the gene.

Perhaps the most dramatic result of this study was the effectiveness of *α-dsrB* and *α-bolA* in restoring carbapenem susceptibility to *E. coli* CUS2B. At 10 µM, each PNA showed significant synergistic interaction with sub-MIC ertapenem, and *α-bolA* showed significant synergistic interaction with sub-MIC meropenem at a concentration of 15 µM. Surprisingly, each *S*-value for these three combinations (0.85, 0.83, and 0.61) was comparable to the value of *α-ampC* (*S* = 0.76). Each of these results is consistent with the predictions of the RNA sequencing: *dsrB* was found to be overexpressed at both time points for ertapenem, but only at 30 minutes for meropenem, and *bolA* was found to be overexpressed in both at 60 minutes. Although little is known about *dsrB*, it is reported to be regulated by the general stress response sigma factor RpoS.^60^ Inhibition of DsrB translation abolished bacterial growth in combination with ertapenem (*S* = 0.85, Fig. 4), which is consistent with previous research that has associated β-lactam antibiotics with induction of the RpoS general stress response regulon. Importantly, though, our results confirm that activation of DsrB is specific to the transcriptome of a resistance phenotype, and that its upregulation is a necessary component of the CRE response in this clinical isolate. BolA is known to be induced under a variety of stress conditions^59^ and is involved in biofilm induction^81–83^ and protective morphological changes^58^ for *E. coli* cells. Perhaps most important, though, is that BolA has been shown to control expression of the proteins PBP5 and PBP6 in *E. coli*, as well as the β-lactamase AmpC.^58^ Although neither *E. coli* PBP5 nor PBP6 has been shown to bind meropenem or ertapenem—these antibiotics are known to bind preferentially to PBP2 and PBP3^18, 24, 41, 72^—our previous results indicate that AmpC contributes to the observed ertapenem-resistance phenotype. The effect of *α-bolA* differs from that of *α-ampC*, however, in its effect on meropenem efficacy. This finding provides strong evidence that BolA is important to the general carbapenem response in *E. coli*, and is not merely incidentally upregulated in the meropenem response, as might be concluded from transcriptomic analysis alone. Additionally, the success of both the *α-dsrB* and *α-bolA* in PNA-carbapenem combination treatments—in three experiments fully abolishing growth (Fig. 4)—represents the first time that transcriptomic analysis has been used to design PNA antibiotics, and, importantly, shows equal if not better efficacy than the genome-derived PNA *α-ampC*. Further development of this strategy will aid in the design of antisense inhibitors to target pathogenic species about which less information—essential genes, genome annotation, etc.—is readily available.

The final PNA for which we find interaction with carbapenem treatment is *α-ygaC*. Similar to *α-flhC*, this PNA showed the ability to rescue growth in *E. coli* CUS2B in cultures treated with previously lethal meropenem concentrations, but did not replicate this effect in ertapenem-treated cultures. Though we observed overexpression of the unannotated RNA transcript antisense to *ygaC*, not the gene itself, the PNA demonstrated surprising effectiveness at inducing resistance. These results affirm that the *ygaC* antisense transcript is likely controlling the gene’s expression, and suggest that *ygaC* downregulation is associated with a more broadly resistant phenotype. The growth rescue in meropenem-treated conditions by both *α-ygaC* and *α-flhC* points to routes that resistant bacteria may take towards developing broad resistance. Furthermore, the success of *α-ygaC*, considering that it is a largely unstudied gene never before linked to antibiotic resistance, demonstrates the specific utility of transcriptomics in the design of PNA for antibiotic applications.

Resistance has been extensively studied at the genetic level, which has helped researchers to understand many genetic factors that can contribute to resistance. However, carbapenem resistance is not often studied at a transcriptomic level, and, when it is, the research almost exclusively examines basal expression levels.^30–35^ In this work, we profiled the short-term transient transcriptomic responses of a partially carbapenem-resistant clinical isolate of *E. coli* and used our FAST platform to design PNA that uncovered the importance of multiple systems in the development of this phenotype, including the regulators *flhC*, *hycA*, and *bolA*. Furthermore, by inhibiting *hycA*, *bolA*, and *dsrB* with FAST PNA, we were able to re-sensitize the clinical isolate to sub-MIC carbapenem concentrations. There are no small molecules known to inhibit the action of any of these gene targets, which demonstrates the value of PNA inhibitors to fill in the gaps in modern antibiotic discovery. This is the first time that transcriptomic analysis has been applied to identify targets for antibiotic PNA applications, and it is striking how effective the transcriptome-derived PNA were in inducing carbapenem susceptibility to the CRE isolate. We find two such PNA-carbapenem combinations that are more effective than the anti-β-lactamase PNA *α-ampC*. In future research we will seek to develop these results into a means to consistently and systematically re-sensitize MDR pathogens to carbapenems, and thus maintain the efficacy of these antibiotics of last resort.

## Methods

### Strain and culture conditions

*E. coli* CUS2B was provided the Dr. Nancy Madinger, at the University of Colorado Hospital Clinical Microbiology Laboratory’s organism bank. The isolate was obtained via rectal swab from a 29-year old pregnant female patient. Unless otherwise mentioned, CUS2B was propagated in aerobic conditions at 37°C in liquid cultures of cation-adjusted Mueller Hinton broth (CAMHB) with shaking at 225 rpm or on solid CAMHB with 15 g/L of agar.

### Minimum inhibitory concentration assays

Three colonies were picked from a plate and used to inoculate three separate overnight cultures in 1 mL CAMHB each. After 16 hours, the cultures were diluted 1:10,000 and treated with a range of antibiotic concentrations, decreasing in 2-fold increments, in a 384 or 96-well microplate using three biological replicates per condition. Growth in the plate was monitored with a Tecan GENios (Tecan Group Ltd.) running Magellan™ software (v 7.2) at an absorbance of 590 nm every 20 minutes for 16 hours, with shaking between measurements. The minimum inhibitory concentration was identified as the lowest antibiotic concentration preventing growth. *Genome sequencing*

Five colonies were picked from a plate and resuspended in liquid culture. After 16 hours, 1 mL of culture was used for genomic DNA isolation with the Wizard DNA Purification Kit (Promega). Approximately 2 µg of DNA was used to prepare a paired-end 250-bp sequencing library with the Nextera XT DNA library kit. The library was sequenced on an Illumina MiSeq, resulting in 407,910 reads (20X coverage). The *de novo* assembly is 5,325,941 bp in length with a GC content of 50.59%. The largest contig is 394,969 bp, and the N50 is 100,215. The genome contains 5,360 protein coding sequences, 114 RNA coding sequences, 82 tRNAs, 11 ncRNAs, 260 pseudogenes, and 2 CRISPR arrays.

The FASTQ files were filtered for quality using Trimmomatic (v0.32)^84^, in sliding window mode with a window size of 4 bases, a minimum average window quality of 15 (phred 33 quality score), and a read length of at least 36 bases. For resequencing, reads were aligned to various *E. coli* RefSeq reference genomes using Bowtie 2 (v2.2.3)^85^. SAMTools (v0.1.19)^86^ was used to remove PCR duplicates and create indexed, sorted BAM files. Variants were called and filtered using the Genome Analysis Toolkit (v2.4-9).^87^ To pass the filter, a SNP had to meet the following criteria: QD < 2.0, FS > 60, MQ < 40.0, ReadPosRankSum < −2.0, and MappingQualityRankSum < −12.5. Filter criteria for indels was: QD < 2.0, FS > 60.0, and ReadPosRankSum < −2.0. A custom Python script was used to annotate variant call files using the corresponding GFF from RefSeq. For *de novo* assembly, reads were assembled using SPAdes (v 3.5.0)^88^ and annotation was performed with the NCBI Prokaryotic Genome Annotation Pipeline (v 4.0)^89^. Quality of the assembly was assessed using QUAST^90^. The FASTA generated by SPAdes was used for MLST^91^, identification of resistance genes with ARG-ANNOT^39^, locating CRISRRs with CRISPRfinder^92^, and locating plasmids with PlasmidFinder^93^.

### RNA-sequencing and differential expression analysis

Colonies were picked from a plate and resuspended in liquid culture. After 16 hours of growth, the culture was diluted 1:20 into duplicate 15 mL cultures. These were grown for 1 hour, then 3 mL from each was preserved in 2 volumes of RNAprotect. Each culture was divided into three equal parts. No antibiotic was added to one part and antibiotic were added to the other two for final concentrations of 2 μg/mL of ertapenem or 1 μg/mL of meropenem, corresponding to 50% of the MIC under these growth conditions (note that these conditions are different than the procedures used to determine the MICs in Fig. 1). After thirty and sixty minutes of growth, 1.5 mL of culture was collected from each and stored in RNAprotect. Cultures were flash frozen in ethanol and dry ice and stored at −80⁰C until the time of RNA extraction.

To extract RNA, samples were thawed and resuspended in 100 μL TE buffer with 0.4 mg/mL lysozyme and proteinase K. After incubation at room temperature for 5 minutes, 300 μL of lysis buffer with 20 μL/mL β-mercaptoethanol was added to each and vortexed to mix. Each lysis solution was split in half, with one half being processed for total RNA isolation and the other for small RNA isolation. Total RNA was isolated using the GeneJet RNA Purification kit (Thermo Scientific) followed by DNase treatment with the TURBO DNA-free kit (Ambion). Small RNA was isolated using the *mir*Vana miRNA isolation kit (Thermo Scientific). Concentration and A260/A280 were measured on a Nanodrop 2000 (Thermo Scientific). A minimum of 130 ng of RNA per sample was submitted for sequencing library preparation. Quality was further assessed with a Bioanalyzer (Agilent). Libraries were prepared using the RNAtag-Seq protocol,^94^ wherein individual samples are barcoded and pooled prior to ribosomal RNA treatment and cDNA synthesis. Here, the total RNA samples were combined into one pool, and the small RNA enriched samples were combined into a separate pool. The total RNA pool was fragmented via incubation with FastAP buffer at 94°C. After another DNAse treatment, barcoded adapters were ligated with T4 DNA ligase, then the samples were pooled and subjected to Ribo-Zero treatment. The prep for small RNA libraries was similar, without the fragmentation or ribosomal RNA treatment steps. Total RNA libraries were sequenced on a NextSeq 500 (Illumina) using a high output cycle 75-cycle run. Small RNA libraries were sequenced on a NextSeq with a medium output run, halted after 75 cycles.

FASTQ files were demultiplexed with the barcode splitter function from the FastX toolbox (http://hannonlab.cshl.edu/fastx_toolkit/, v0.0.13.2). The first seven base calls (containing the barcode) were trimmed using FastX, then adapters were removed and all reads were trimmed for quality using the sliding window mode in Trimmomatic (v0.32)^84^. Quality of the resulting FASTQ files was validated with FastQC (http://www.bioinformatics.babraham.ac.uk/projects/fastqc/). For total RNA data (including sense and antisense transcripts) and differential expression analysis, reads were aligned to the draft assembly of the CUS2B genome (available as RefSeq GCF_001910475.1) with Bowtie 2 (v2.2.3). An average of 16.5±3.2 million reads were successfully mapped per sample. SAMTools (v0.1.19) was used to create bam files. HTSeq (v0.6.1)^95^ was used to build count tables for sense and antisense reads. DESeq^47^ was used to determine differentially expressed genes, with a pooled dispersion metric and a parametric fit. Genes were considered significantly differentially expressed if the Benjamini-Hochberg adjusted *P*-value was less than 0.05. For small RNA data, trimmed FASTQ files were submitted to Rockhopper,^57^ which mapped to the *E. coli* UMN026 genome (90,152±25,642 reads mapped per sample) and performed differential expression analysis. Small RNA transcripts were considered significantly DE if the Benjamini-Hochberg adjusted *P*-value was less than 0.05.

### PNA design

PNA design was carried out using the PNA Finder toolbox.^36^ The toolbox is built using Python 2.7, the alignment program Bowtie 2,^85^ the read alignment processing program SAMtools,^86^ and the feature analysis program BEDTools.^96^ The toolbox takes a user-provided list of gene IDs and cross-references the IDs against a genome annotation file to determine the feature coordinates for each ID. The toolbox then uses these coordinates to extract PNA target sequences of a user-specified length (12 bases in this study) and user-specified positions relative to the start codon from a genome assembly FASTA file. PNA Finder provides a list of PNA candidates (the reverse complements of the target sequences) and sequence warnings regarding solubility and self-complementation. Finally, PNA Finder screens the list of PNA candidates against a user-provided genome assembly (in this study, the genome for *E. coli* CUS2B) to search for off-targets, and uses this analysis to filter the candidates into a final PNA list for synthesis.

### PNA synthesis

PNA were synthesized using an Apex 396 peptide synthesizer (AAPPTec, LLC) with solid-phase Fmoc chemistry at a 10 µmol scale on MBHA rink amide resin. Fmoc-PNA monomers were obtained from PolyOrg Inc., with A, C, and G monomers protected with Bhoc groups. PNA were synthesized with the N-terminal cell-penetrating peptide (KFF)_3_K. Cell-penetrating peptide Fmoc monomers were obtained from AAPPTec, LLC, and lysine monomers were protected with Boc groups. PNA products were precipitated in diethyl ether and purified as trifluoroacetic acid salts via reverse-phase HPLC using a C18 column.

### PNA-antibiotic interaction assays

Three colonies were picked from a plate and used to inoculate three separate overnight cultures in 1 mL CAMHB each. After 16 hours, the culture was diluted 1:10,000 in a 384-well microplate using three biological replicates per condition. The total culture volume for each treatment was 50 µL. PNA were stored at −20 °C dissolved in 5% v/v DMSO in water. Growth in the plate was monitored with a Tecan GENios (Tecan Group Ltd.) running Magellan™ software (v 7.2) at an absorbance of 590 nm every 20 minutes for 24 hours, with shaking between measurements.

Interaction effects were evaluated for significance using a two-way ANOVA test, and *S*-values were calculated with respect for the expected growth inhibition as calculated by the Bliss Independence model.^66^ The *S*-value for a given timepoint was calculated as follows:

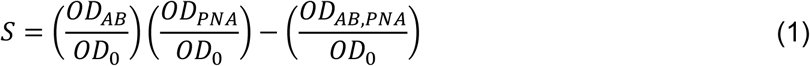

For a given timepoint (24 hours was used in our analyses), the variable OD_AB_ represents optical density with only carbapenem treatment, OD_0_ represents the optical density without treatment, OD_PNA_ represents optical density with only antisense-PNA treatment, and OD_AB, PNA_ represents the optical density with a combination treatment. Plus/minus for *S*-values (Table 1) was calculated by propagating standard error values for each term in Equation 1.

### Other software and resources utilized

The clustergram function from MATLAB’s Bioinformatics toolbox was used for building heatmaps and dendograms. A Euclidean distance metric, optimal leaf ordering, and average linkage function were used for clustering. Ecocyc^97^ was used to gain gene names and descriptions and to define functional classes. NCBI’s BLAST^98^ was used to predict gene function and to determine similarity of sequences in CUS2B to other bacterial strains. PANTHER^50^ was used for statistical overrepresentation tests, with a Bonferroni correction applied to all reported *P*-values.

### Data availability

The whole genome shotgun sequencing data has been deposited at DDBJ/ENA/GenBank under the accession MSDR00000000.1. The version described in this paper is version MSDR01000000. The RNA sequencing data has been deposited in NCBI’s Sequence Read Archive under the accession SRP101716.

## Supporting information

Supplemental small RNA Data

Supplemental total RNA Data

## Author contributions

K.E.E., T.R.A., and A.C. conceived of the study.. K.E.E. and T.R.A. performed all experiments and analysis. K.E.E., T.R.A., and A.C. wrote the manuscript.

## Acknowledgments

This work was funded by University of Colorado Dean’s Graduate Research Grants (K.E.E. and T.R.A.), the University of Colorado GAANN Fellowship (T.R.A.), the W.M. Keck Foundation, the DARPA Young Faculty Award (D17AP00024) (A.C.) and the National Aeronautics and Space Administration (NASA) Cooperative Agreement Notice – Translational Research Institute (TRISH) (NNX16A069A) (A.C.). We thank the University of Colorado BioFrontiers Institute Next-Gen Sequencing Core Facility, which performed all Illumina sequencing and library construction.

