## Supplemental Information for "Transcriptome-based design of antisense inhibitors re-sensitizes CRE *E. coli* to carbapenems"

#### Supplemental methods

*Sequence typing*

*E. coli* do not cluster cleanly into phylogenic trees. For instance, non-pathogenic and pathogenic strains appear in each of the four main groups (A, B1, B2, and D), and even the specific pathogenic type, such as enterohaemorrhagic (EHEC), enteropathogenic (EPEC), and enteroinvasive (EIEC) is dispersed among phylogenetic groups.[1] Heterogeneity among phylogenetic groups has been speculated to be due to high levels of recombination or to rapid radiation after bottlenecking.[1] Nevertheless, information useful in characterization can be obtained by determining sequence type and phylogenetic grouping. We used the draft genome assembly to classify *E. coli* CUS2B per the MultiLocus Sequencing Typing (MLST 1.8) tool from the Center for Genomic Epidemiology.[2] We employed the MLST from Jaureguy et. al.,[3] which determines type using the sequence of eight housekeeping genes. By this scheme, the isolate was found to match ST-44 in phylogenetic group D. Group D has found to have higher mutation rates, as a whole, when compared to other groups, but is also a very diverse group, with members including K1, EHEC, EIEC, Shigella, “other” pathogens, and non-pathogenic strains.[1]

*Analysis of mutations contributing to fluoroquinolone, rifampicin, or nitrofurantoin resistance*

We did not identify any ARG that could impart fluoroquinolone, rifampicin, or nitrofurantoin resistance, so we searched the genome sequence for mutations known to convey resistance to these antibiotics (Table S4). Comparing the sequence of CUS2B to that of the antibiotic-susceptible strain *E. coli* MG1655, we find codon changes at positions 83 and 87 in the DNA gyrase subunit A gene *gyrA*; mutations in these positions have been previously revealed to result in fluoroquinolone resistance[4]. There were four codon changes in DNA gyrase subunit B (*gyrB*). None of the four positions fell in a previously determined resistance-determining region,[5] though it has been suggested that *gyrA* mutations are more frequent and bestow a selective advantage for fluoroquinolone resistance in *E. coli.*[6] There were five codon changes identified in *nfsA* and three in *nfsB*, NADPH nitroreductase genes that are nitrofurantoin resistance determinants. None of the eight positions matched those from previous studies analyzing the genetic basis for nitrofurantoin resistance;[7,8] however, the wide range of resistance-conferring mutations previously identified suggests that there may not be a narrow resistance-determining region in *nfsA/B*. We did not find any nonsynonymous mutations in *rpoB*, which is a determinant of rifampicin resistance.[9] Thus, the low-level rifampicin resistance observed may be a result of either a non-specific antibiotic response, or mutations in other genes not commonly linked to rifampicin resistance.

*Antisense RNA involved in carbapenem-resistance*

In this analysis, we located antisense RNA that were DE in response to ertapenem and meropenem treatment and summarized the potential functions that these RNAs could influence via antisense binding (Fig. S4). The percentage of differentially expressed antisense transcripts ranged between 4% (in ertapenem at 60 minutes) and 23% (in meropenem at 60 minutes) of the total number of transcripts (Fig. S4A). More antisense transcripts were significantly overexpressed than underexpressed in all cases, and no underexpressed antisense transcripts were identified in either ertapenem or meropenem at 60 minutes. We grouped the antisense transcripts identified in ertapenem or meropenem by the biological function of the sense gene (Fig. S4B). The plurality of these corresponding sense genes did not have a known function (31% in ertapenem, 47% in meropenem). Of the genes with known function, we find functions related to transport, metabolism, oxidation-reduction, protease, and transcriptional regulation. Functions unique to overexpressed transcripts in ertapenem include amino acid processing, DNA processing, membrane, signal transduction, and stress response. Two antisense transcripts were DE only in ertapenem, at both 30 and 60 min (Fig. S4C). *psiE* is associated with stress response, while the BTW13_RS28340 is of unknown function, and each was overexpressed with respect to the no treatment condition. Two other transcripts were significantly overexpressed in ertapenem at 30 and 60 minutes, in meropenem at 30 minutes, but not in meropenem at 60 minutes. One of these is antisense to the regulator *zur* and the other, BTW13_RS03075, is of unknown function (Fig. S4C).

### Supplemental Tables and Figures

**Table S1** Comparison of four genes related to β-lactam resistance located in the genome of CUS2B with homologues in *E. coli* strains MG1655 and UMN026. Codon changes are bold if they were identified in comparison with both MG1655 and UMN026. Genes were identified using the ARG-ANNOT database.

| Gene in CUS2B | Homologue enzyme type | *E. coli* MG1655 homologue | Mutations vs MG1655 | *E. coli* UMN026 homologue | Mutations vs UMN026 |
|---|---|---|---|---|---|
| *ampC2* | β-lactamase | *ampC* | T4M, **S102I**, T105A, E140D, **Q196H**, N201T, P209S, N260T, S298I, D367A | *ampC* | **S102I**, **Q196H**, C287W, C325R, T367A |
| *ampC1* | PBP | *yfeW* (PBP4B) | V53A, S57G, **N272D**, T388A | hypothetical protein | T23P, A141T, **N272D**, S327N |
| *ampH* | PBP | *ampH* | **M1L**, V83I | *ampH* | **M1L**, Y41H |
| *Generic PBP* | PBP | *mrdA* (PBP2) | none | *mrdA* | none |

**Table S2** Differentially expressed genes from select total RNA-seq overlap categories. ERT = ertapenem, MER = meropenem.

| ERT 30 minutes and MER 30 minutes (41 genes) | ERT 30 minutes and ERT 60 minutes (38 genes) |
|---|---|
| *alaE* | *[citrate (pro-3S)-lyase] ligase* |
| *bdm* | *4-alpha-glucanotransferase* |
| *bifunctional N-acetylglucosamine-1-phosphate uridyltransferase/glucosamine-1-phosphate acetyltransferase* | *50S ribosomal protein L11* |
| *EEP domain-containing protein* | *alpha-glycosidase* |
| *flagellar L-ring protein* | *carbamoyl-phosphate synthase large chain* |
| *flgA* | *carbon starvation protein A* |
| *flgC* | *citrate (pro-3S)-lyase subunit beta* |
| *flgD* | *citrate lyase ACP* |
| *flgE* | *citrate lyase subunit alpha* |
| *flgF* | *degP* |
| *flgG* | *dihydroorotate dehydrogenase (quinone)* |
| *flgI* | *emrA* |
| *flgJ* | *formate hydrogenlyase complex iron-sulfur subunit* |
| *flgK* | *formate hydrogenlyase subunit 3* |
| *flhA* | *hycA* |
| *flhB* | *hycD* |
| *fliA* | *hycE* |
| *fliD* | *hydN* |
| *fliF* | *hypA* |
| *fliG* | *hypothetical protein (2)* |
| *fliH* | *lamB* |
| *fliI* | *loiP* |
| *fliJ* | *lptG* |
| *fliK* | *malE* |
| *fliL* | *malF* |
| *fliM* | *malG* |
| *fliN* | *malK* |
| *fliO* | *maltodextrin phosphorylase* |
| *fliP* | *membrane protein* |
| *fliZ* | *NADH dehydrogenase* |
| *glutamine--fructose-6-phosphate aminotransferase* | *phage shock protein C* |
| *hypothetical protein (5)* | *phage shock protein G* |
| *IlvGMEDA operon leader peptide* | *phoH* |
| *N-acetylmuramic acid 6-phosphate etherase* | *plaP* |
| *pyruvate dehydrogenase complex repressor* | *psiE* |
| *stress-induced protein, UPF0337 family* | *translation initiation factor* |
| *transketolase* | |

**Table S3** PNA targets and antisense PNA sequences, start codons in bold

| Gene Target in *E. coli* CUS2B | Target Sequence (5’ to 3’) | PNA Sequence (N to C) |
|---|---|---|
| *ampC* | CAGACCCT**ATG**T | $(KFF)_3$K-(AEEA)-A**CAT**AGGGTCTG |
| *bolA* | TCATG**ATG**ATAC | $(KFF)_3$K-(AEEA)-GTAT**CAT**CATGA |
| *dsrB* | **CTG**CCAGAAAAA | $(KFF)_3$K-(AEEA)-TTTTTCTGG**CAG** |
| *flhC* | TGGGAATA**ATG**C | $(KFF)_3$K-(AEEA)-G**CAT**TATTCCCA |
| *hycA* | TGACA**ATG**ACTA | $(KFF)_3$K-(AEEA)-TAGT**CAT**TGTCA |
| *malT* | GTGATTAACT**AT** | $(KFF)_3$K-(AEEA)-**AT**AGTTAATCAC |
| *ygaC* | TCAAAT**ATG**TAT | $(KFF)_3$K-(AEEA)-ATA**CAT**ATTTGA |
| nonsense | ACAGCAAGTGCA | $(KFF)_3$K-(AEEA)-TGCACTTGCTGT |

**Table S4** Codon changes in *E. coli* CUS2B as compared to MG1655 in genes associated with resistance to fluoroquinolones, nitrofurantoins, or rifampicin

| Gene | Nonsynonymous mutations vs MG1655 | Associated resistance |
|---|---|---|
| *gyrA* | S83L, D87N | Fluoroquinolone |
| *gyrB* | A359E, T618A, A653T, V663I | |
| *nfsA* | F9C, D58E, K72Q, T117I, E141K | Nitrofurantoin |
| *nfsB* | D66G, V93A, E137Q | |
| *rpoB* | none | Rifampicin |

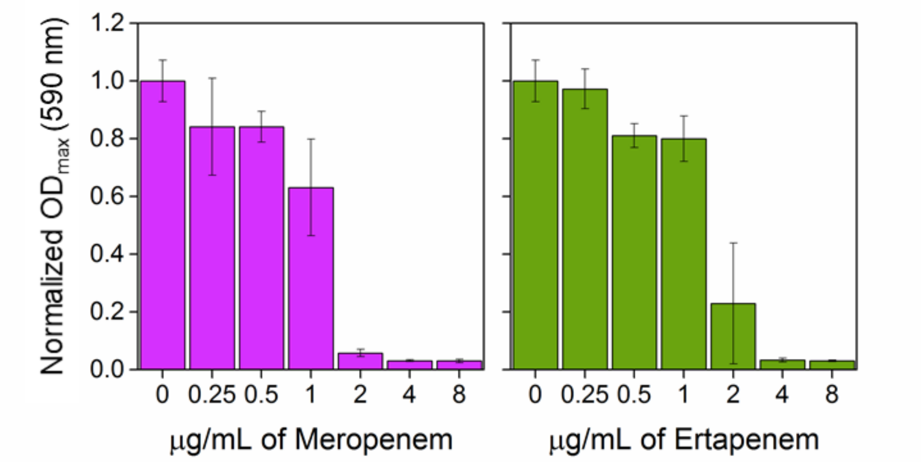


**Figure S1** Maximum optical density for each antibiotic concentration, normalized to an untreated control. Growth conditions here are as in the RNA-seq experiment: a 1:20 dilution from an overnight culture was grown for 1 hour before the addition of antibiotics.

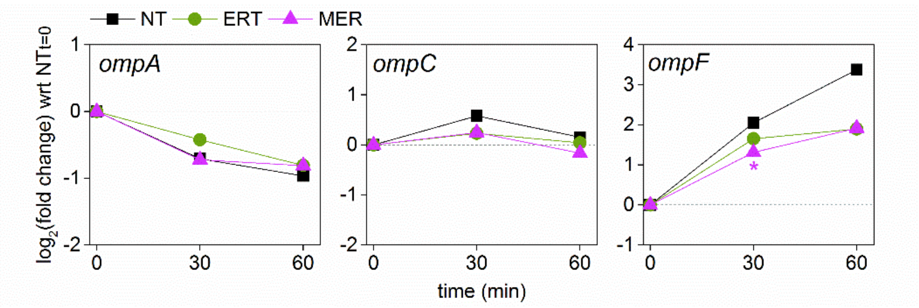


**Figure S2** Time course of gene expression for the outer membrane porin genes *ompA*, *ompC*, and *ompF*. An asterisk is used to indicate a significant differential expression change ($P < 0.05$) vs the no treatment condition at the same timepoint. NT = no treatment, ERT = ertapenem, MER = meropenem

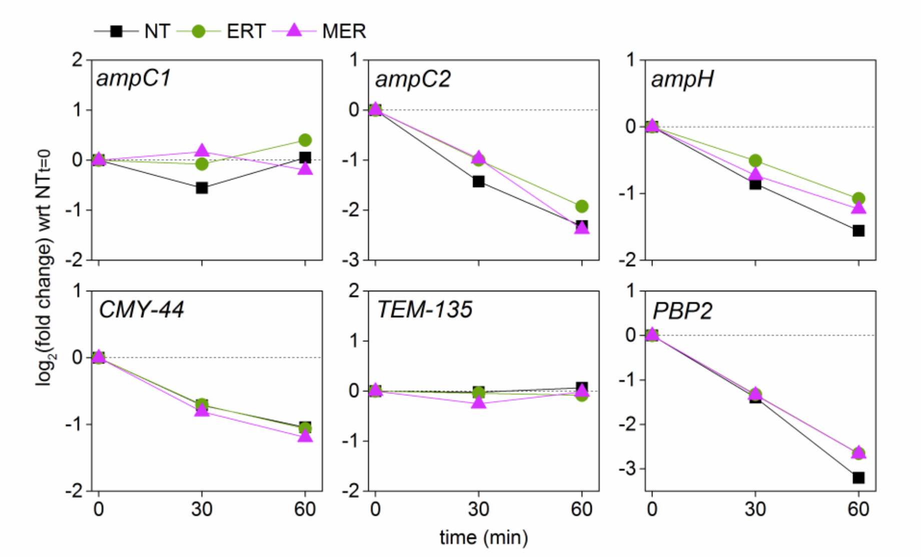


**Figure S3** Time course of gene expression for the β-lactam-related genes encoded by *E. coli* CUS2B. None of these genes were significantly differentially expressed with respect to the no treatment samples at the same timepoint. NT = no treatment, ERT = ertapenem, MER = meropenem.

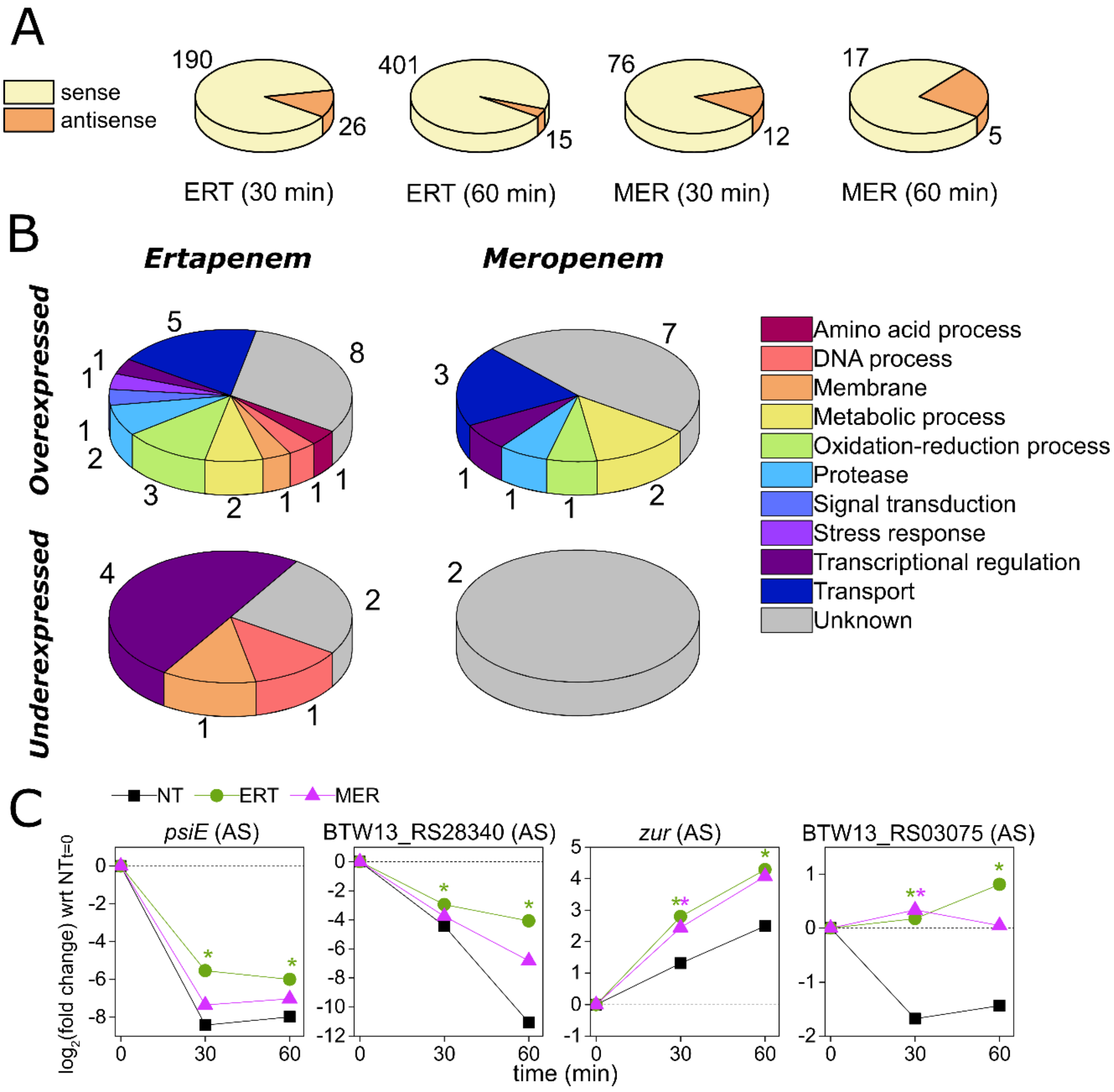


**Figure S4** (A) Percentage of differentially expressed sense and antisense transcripts in each experimental condition. Number of transcripts in each case is indicated. (B) Percentage of differentially expressed antisense transcripts, by functional class. Transcripts from 30 and 60 minutes are grouped together for this analysis, within their respective antibiotic treatment category. (C) Time course of four antisense transcripts of interest. Log2(fold change) here is with respect to the no treatment condition at time t=0. NT = no treatment, ERT = ertapenem, MER = meropenem

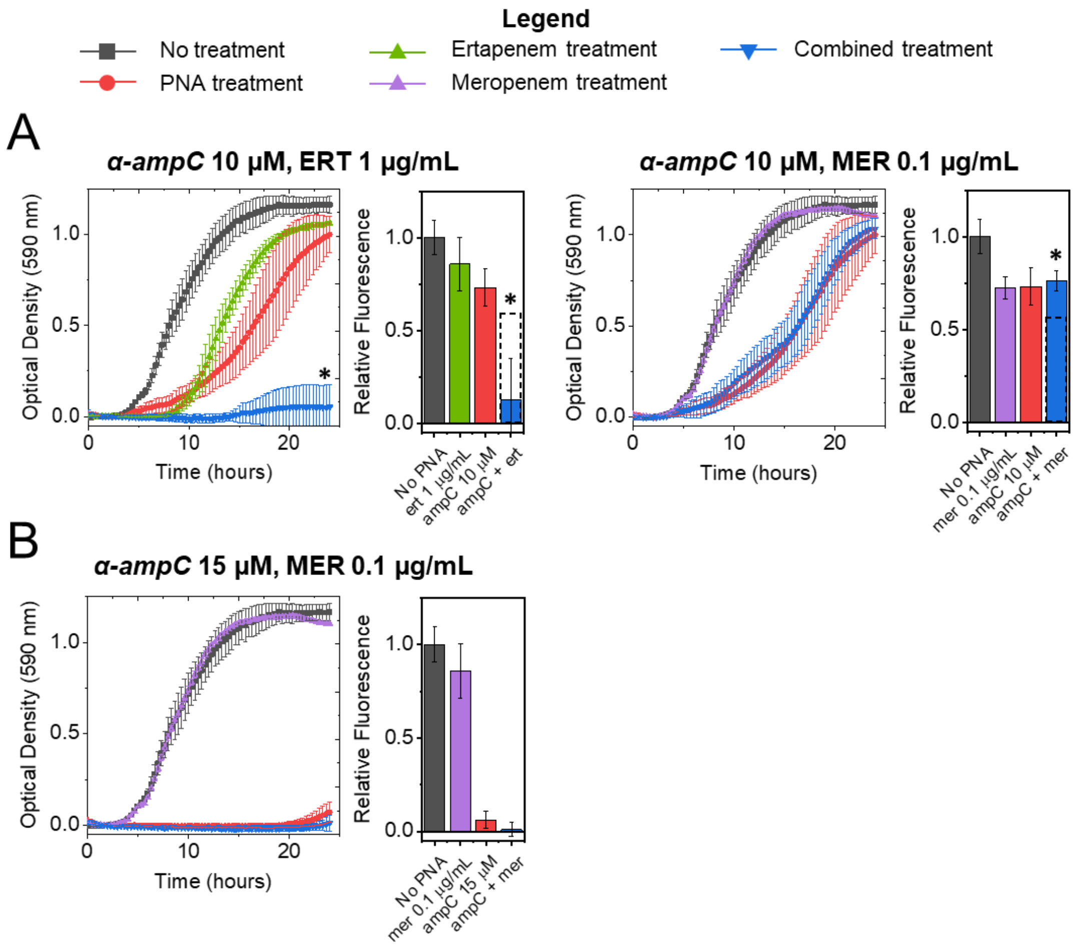


**Figure S5** Growth curves and cell viability assays for *E. coli* CUS2B treated with a combination or ertapenem or meropenem. (A) α-*ampC* at 10 μM combined with ertapenem at 1 μg/mL and meropenem at 0.1 μg/mL. Significant synergistic interaction was observed in the growth curve endpoint and viability assay for α-*ampC* and ertapenem. Significant antagonistic interaction interaction was observed in the α-*ampC* at 10 μM and meropenem combination, though it was not found in the growth curve endpoints. (B) (A) α-*ampC* at 15 μM combined with meropenem at 0.1 μg/mL. No significant interaction was observed.

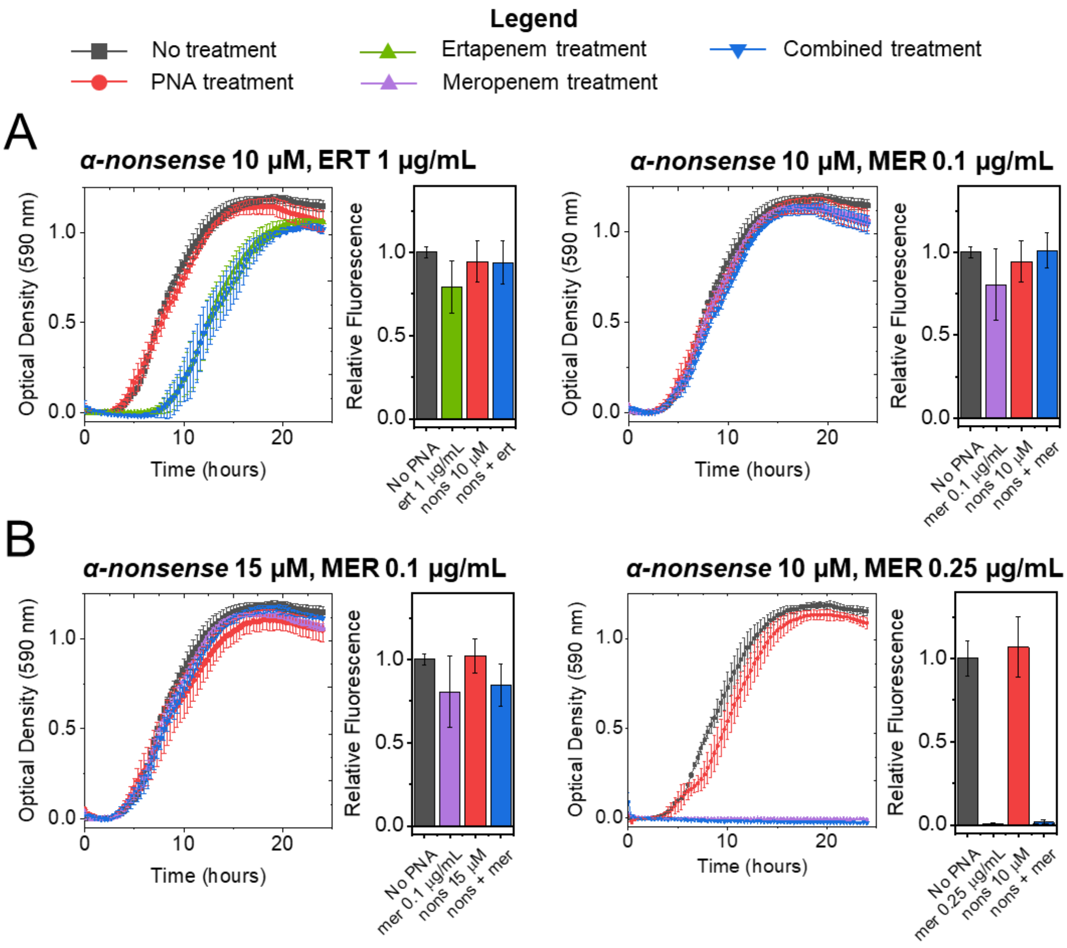


**Figure S6** Control growth curves and endpoint viability assays for *E. coli* CUS2B treated with a combination of a scrambled nonsense PNA and each carbapenem antibiotic, shown for each condition in which significant interaction was found between an antisense inhibitory PNA and carbapenem antibiotic. (A) Initial PNA trial concentrations: *α-nonsense* at 10 µM combined with ertapenem at 1 µg/mL and meropenem at 0.1 µg/mL. (B) Elevated PNA and antibiotic concentrations: *α-nonsense* at 15 µM combined with meropenem at 0.1 µg/mL and *α-nonsense* at 10 µM combined with meropenem at 0.25 µg/mL.

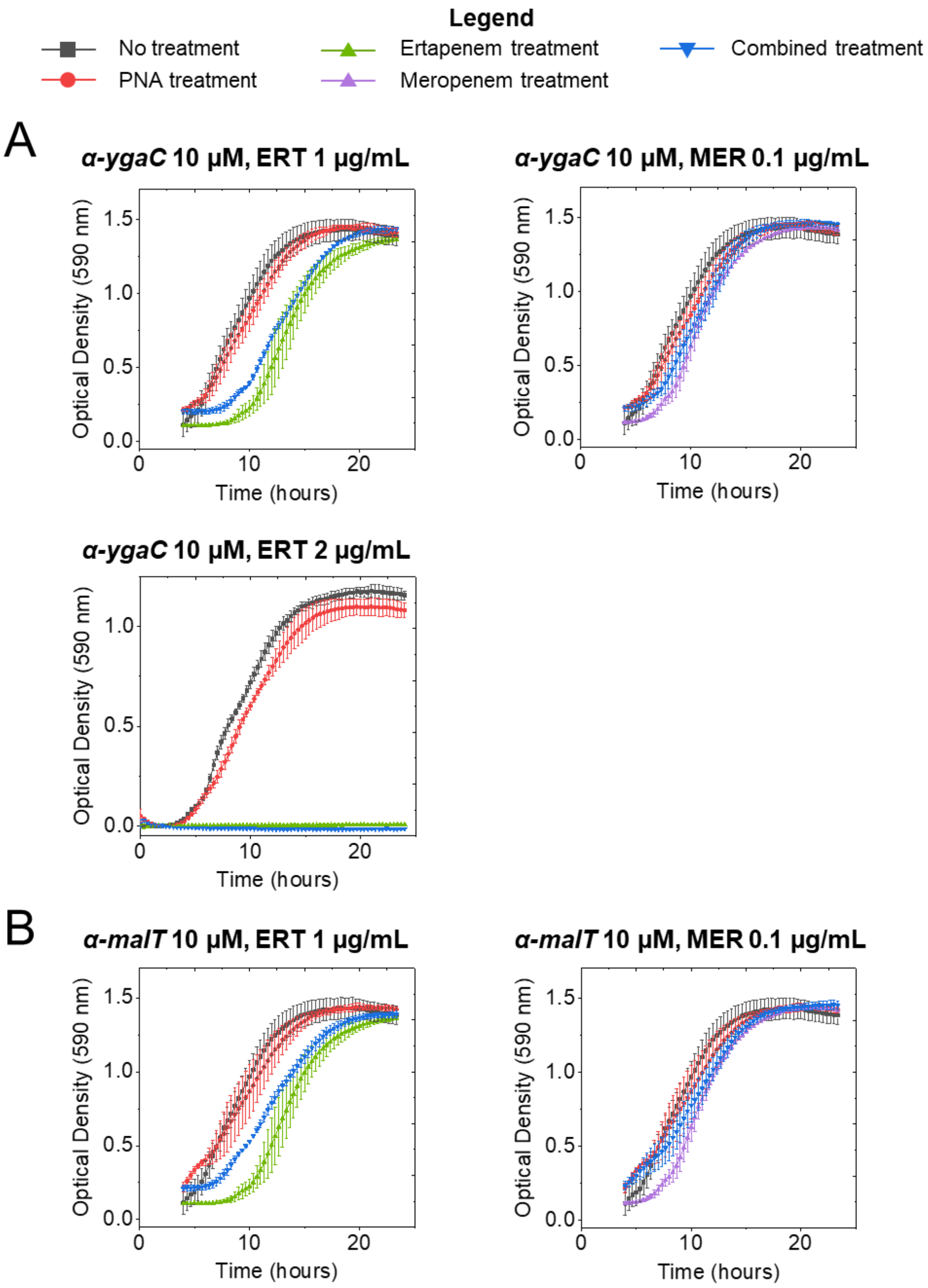


**Figure S7** Growth curves for *E. coli* CUS2B treated with a combination of (A) α-*ygaC* or (B) α-*malT* at 10 µM and carbapenem antibiotics (ertapenem at 1 µg/mL or meropenem at 0.1 µg/mL). None of the experiments displayed evidence of interaction between the PNA and antibiotic treatments, and these PNA were not pursued further in potentiating the carbapenems. (Note: Though the cultures were incubated with regular shaking for the first 4 hours of treatment, measurement data was lost. As such, growth curves are normalized by subtracting non-culture microplate wells containing broth, instead of subtracting early timepoints as with the main text data).

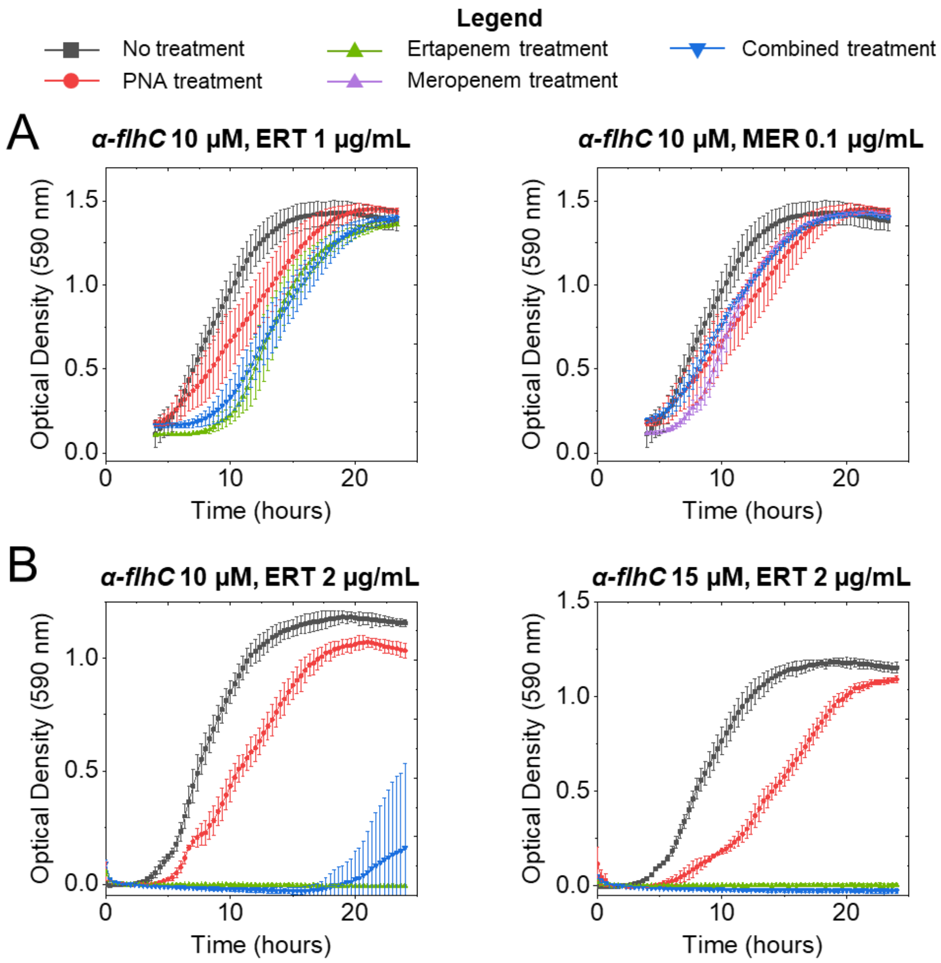


**Figure S8** (A) Growth curves for the α-*flhC*/carbapenem combinations at sub-MIC carbapenem concentrations. (B) Growth curves for the α-*flhC*/MIC ertapenem treatment growth rescue experiments. None of the experiments displayed evidence of interaction between the PNA and antibiotic treatments. (Note: Though the cultures for the top two panels were incubated with regular shaking for the first 4 hours of treatment, measurement data was lost. As such, growth curves are normalized by subtracting non-culture microplate wells containing broth, instead of subtracting early timepoints as with the main text data).

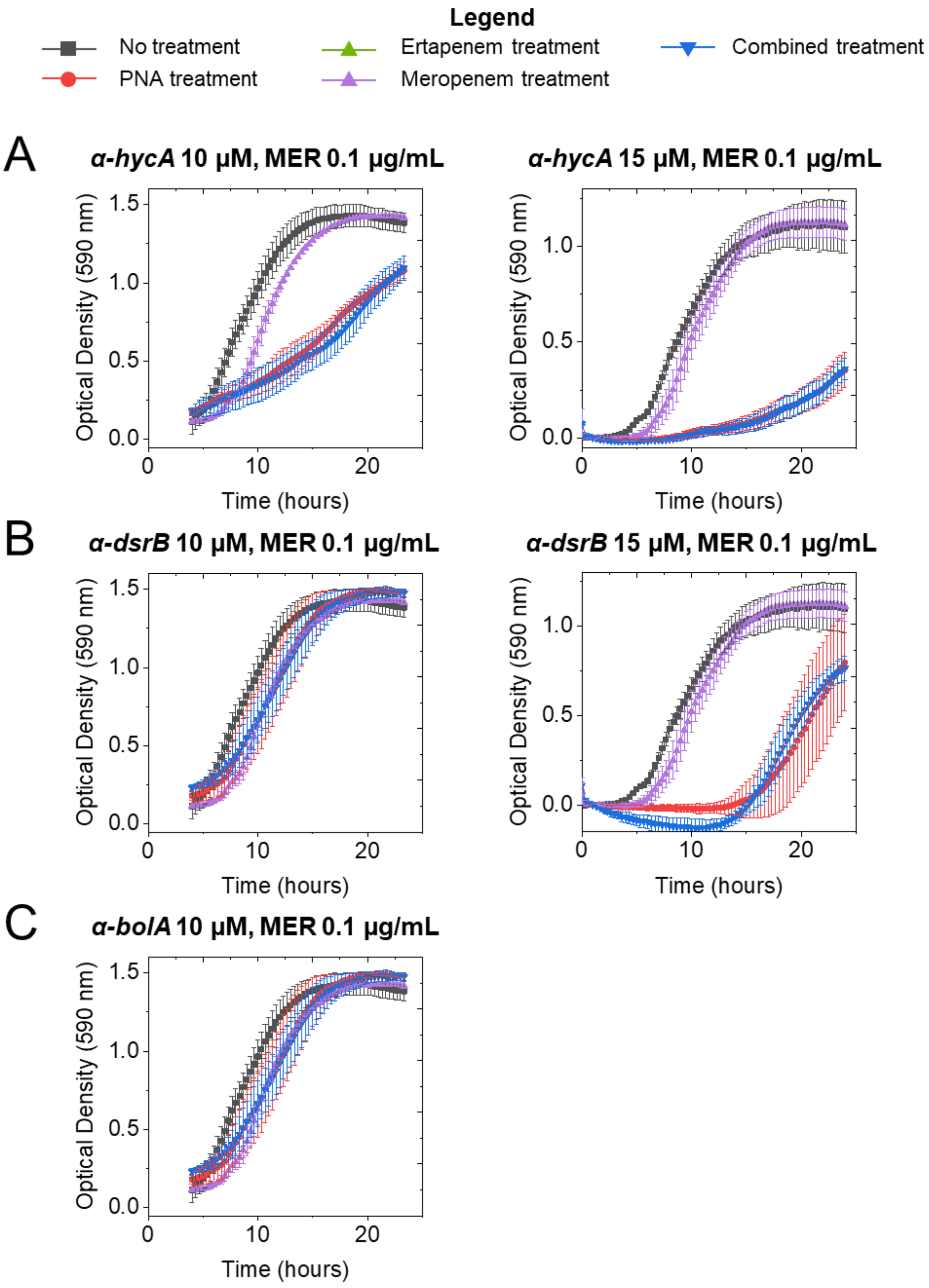


**Figure S9** Growth curves for *E. coli* CUS2B treated with a combination of (A) α-*hycA*, (B) α-*dsrB*, or (C) α-*bolA* at 10 µM and 15 µM, and meropenem at 0.1 µg/mL. None of the experiments displayed evidence of interaction between the PNA and antibiotic treatments. (Note: Though the cultures for the left-side panels were incubated with regular shaking for the first 4 hours of treatment, measurement data was lost. As such, growth curves are normalized by subtracting non-culture microplate wells containing broth, instead of subtracting early timepoints as with the main text data).
